# Immunological Differences in Atopic Dermatitis Across Age Groups: Insights from Single-Cell Multi-Omics

**DOI:** 10.1101/2025.11.22.689843

**Authors:** Gian Carlo L. Baldonado, Sugandh Kumar, Joy Jin, Xiaohui Fang, Alexander Ildardashty, Mitchell Braun, Isaac M. Neuhaus, Erin Mathes, Tina Bhutani, Wilson Liao

## Abstract

**Background:** Atopic dermatitis (AD) occurs across all ages but presents distinct clinical and immunologic features between children, adults, and older adults. The molecular programs underlying these age-specific immune differences remain poorly understood.

**Methods:** We performed single-cell multi-omics profiling of peripheral blood mononuclear cells (PBMCs) from 29 AD patients and 29 matched healthy controls (HC), spanning pediatric (0–17 years), adult (18–59 years), and geriatric (≥60 years) groups. Using Cellular Indexing of Transcriptomes and Epitopes by sequencing (CITE-seq), we simultaneously quantified transcriptomic (RNA) and surface proteomic (ADT) profiles across ∼280,000 immune cells. Integrated analyses identified 30 immune subsets for cell-type proportion and differential expression analyses. Machine-learning classifiers were trained on significant gene and protein features to distinguish AD subgroups by age.

**Results:** Compared with HC, AD blood showed enrichment of CD14+ monocytes, plasmacytoid dendritic cells, and CD4+ proliferating T cells. Within AD, pediatric patients had increased γδ T cells, naïve CD4+, and naïve CD8+ T cells, while geriatric patients exhibited more CD4+ cytotoxic and CD8+ central memory T cells, indicating a shift from naive to effector predominance with aging. Transcriptomic and proteomic analyses revealed distinct programs: pediatric AD was enriched for IL-10 and cytokine–cytokine receptor signaling; adult AD demonstrated activation of metabolic and nuclear factor kappa-light-chain-enhancer of activated B cells (NF-κB)/Th1/Th17 pathways; and geriatric AD exhibited reduced adaptive immune activity but increased innate signaling. Machine-learning models based on differentially expressed genes and proteins accurately classified AD age groups (transcript-based F1 = 0.70, AUC = 0.79), identifying stable markers such as *IRF2*, *PDK4*, *ZFP90*, CD21, CD94, and CD122.

**Conclusions:** Single-cell multi-omics profiling revealed immune differences across the AD lifespan, transitioning from developmental tolerance in children to inflammatory and metabolic activation in adults to enhanced innate signaling in geriatric individuals. These findings highlight molecular signatures that could support age-specific diagnostics and therapeutic strategies for AD across the lifespan.

## Introduction

Atopic dermatitis (AD) is a chronic, inflammatory skin disease that affects up to 20% of children and 5-10% of adults worldwide^1,2^. Most studies of AD pathogenesis have focused on skin-localized inflammation, revealing epidermal barrier dysfunction, immune polarization, and microbial imbalance as central drivers of disease^3,4,1^. Pediatric AD is typically characterized by T helper 2 (Th2) and T helper 22 (Th22) cytokine dominance, marked by elevated interleukin (IL)-4, IL-13, and IL-31 expression. With age, AD develops stronger T helper 1 (Th1) and T helper 17 (Th17) activity, reflecting broader immune activation and disease chronicity^5,4,6^.

However, AD represents not only a skin barrier-centered disorder but also a systemic immune condition that involves coordinated dysregulation across circulating lymphoid and myeloid compartments^2^. Few studies have investigated peripheral blood mononuclear cells (PBMCs) at single-cell resolution, and age-stratified systemic analyses remain limited. Single-cell peripheral blood profiling of adult AD has revealed expanded interferon- and Janus kinase–signal transducer and activator of transcription (JAK–STAT)-activated monocytes, reduced natural killer (NK)-cell cytotoxicity, and increased co-stimulatory signaling, supporting the concept of systemic immune remodeling^5^. Comparative analyses of pediatric and adult AD have also identified Th2 predominance in children and increased Th1, Th17, and Th22 activity in adults, reflecting age-dependent immune divergence; however, these studies excluded older adults and were limited to transcriptomic data^6^. This gap highlights the need for additional studies examining age-specific assessments of systemic immunity in AD. Here, we performed single-cell multi-omics analysis integrating transcriptomic and proteomic profiling of peripheral blood from pediatric, adult, and geriatric patients and matched healthy controls to delineate age-dependent immune remodeling and identify predictive molecular signatures through integrative machine-learning analysis.

## Materials and Methods

### Subject Recruitment and Sampling

Subjects with atopic dermatitis (AD) were recruited from dermatology clinics at the University of California, San Francisco (UCSF). Healthy controls, matched by age, sex, and race/ethnicity, were recruited at UCSF. A total of 61 AD and healthy subjects were enrolled, stratified across three age groups: pediatric (0–17 years), adult (18–59 years), and geriatric (≥60 years). All AD diagnoses were confirmed by a board-certified dermatologist, and subjects were required to have at least moderate disease severity (validated Investigator Global Assessment for Atopic Dermatitis [vIGA-AD] ≥ 3). Exclusion criteria included current or recent use (within three months) of systemic immunomodulators, active autoimmune disease, active infection, or vaccination within the previous month. Clinical and demographic data collected included age, gender, race/ethnicity, body mass index, and vIGA-AD. Demographic characteristics are given in Supplementary Table 1, Additional File 1. Peripheral venous blood was collected for CITE-seq profiling of PBMCs.

### Sample and Library Preparation

#### Single-Cell Libraries

Peripheral blood was collected in Acid Citrate Dextrose (ACD) Vacutainer tubes and processed within 4 hours of phlebotomy. PBMCs were isolated by Ficoll gradient centrifugation, washed, and cryopreserved in 90% fetal bovine serum (FBS) + 10% dimethyl sulfoxide (DMSO) until use. For CITE-seq, 1×10⁶ cryopreserved PBMCs per subject were thawed, counted on a Vi-CELL XR (Beckman Coulter), and pooled in batches of ten subjects balanced by condition and age group. Following Fc blocking (Human TruStain FcX, BioLegend), cells were stained with a 140-plex TotalSeq-B antibody panel (Supplementary Table 2, Additional File 2) for 30 min at 4°C, washed three times with Cell Staining Buffer, and filtered through 40 µm strainers. Cell viability exceeded 85% in all preparations.

#### Cell Capture and Library Construction

4×10^5^ stained cells were loaded across 8 wells of a 10x Chromium Single Cell 3’ v3.1 chip, targeting approximately 2×10^5^ cell-containing droplets per run. Each droplet encapsulated a single cell for reverse transcription and barcode tagging.

Following lysis, cDNA libraries were prepared according to the manufacturer’s protocol, with separate size selection for gene-expression and antibody-derived tag (ADT) fragments. Libraries underwent amplification using KAPA HiFi HotStart ReadyMix (Kapa Biosystems) and size purification with SPRIselect reagent (Beckman Coulter). Final libraries were quantified by Bioanalyzer 2100 (Agilent) and sequenced on an Illumina NovaSeq 6000 (S4 flow cell) with a target depth of ∼20,000–40,000 reads/cell (RNA) and 5,000–10,000 reads/cell (ADT).

### Genotyping

Genomic DNA was extracted from PBMCs using the DNeasy Blood and Tissue Kit (Qiagen). DNA samples were genotyped using the Affymetrix UK Biobank Axiom Array (ThermoFisher) on a GeneTitan Multi-Channel Instrument (Applied Biosystems). This genotype data was used to enable single cell demultiplexing (see below).

#### Genotype Data Processing

Protocol for genotyping followed Liu et.al^3^. Briefly, single nucleotide polymorphisms (SNPs) with call rate < 95%, Hardy-Weinberg p < 1×10⁻⁵, or minor allele frequency < 5% were removed. Imputation of non-genotyped sites was performed on the Michigan Imputation Server (1000 Genomes Phase 3 v5 Genome Reference Consortium Human build 37 (GRCh37) reference panel; Eagle v2.4 phasing). Next, variant call format (VCF) files were processed using Plink 1.90/2.0, with non-exonic SNPs removed and allele orientation verified using snpflip. Resulting high-quality genotype sets were converted to GRCh38 coordinates using Picard LiftoverVcf. Finally, imputed, quality-filtered genotype data were used as input for Demuxlet to assign each droplet barcode to its subject of origin.

### Single-Cell Data Processing

Raw FASTQ files for RNA and ADT libraries were aligned to the GRCh38 human genome and the ADT feature barcode reference using Cell Ranger v8.0.1 of 10x Genomics with default parameters. Resulting transcript RNA and protein-expression count matrices were further processed by RStudio using Seurat v6. Only droplets identified as non-empty (cell-containing) were retained for downstream analysis.

#### Cell Demultiplexing, Doublet Removal, and Annotation

Demultiplexing was performed using Demuxlet (Popscle suite)^4^, leveraging the imputed SNP genotypes to assign each droplet to its donor and remove inter-individual doublets. Remaining intra-sample doublets were removed using DoubletFinder (v2.0) based on simulated multiplet profiles. For each library, cell-level quality control (QC) metrics, including nCount_RNA, nFeature_RNA, and percent mitochondrial RNA, were evaluated prior to merging.

### Quality Control and Cell Type Annotation

Cells were retained if they met RNA and ADT quality thresholds. RNA-based filtering required nFeature_RNA ≥ 200, nCount_RNA within the interquartile range, and mitochondrial gene content ≤ 15 %. For the ADT modality, cells were retained if total detected protein features were ≤ 260 and if isotype control signals contributed < 2 % of total reads. RNA counts were normalized using SCTransform, regressing out technical covariates including sequencing depth (nCount_RNA) and mitochondrial content (percent.mt). ADT counts were normalized using centered log-ratio (CLR) transformation.

Cell type annotation was performed using the Azimuth reference mapping pipeline integrated in Seurat v6, which projects query datasets onto the Human Cell Atlas PBMC reference (Hao et al., 2021). SCTransform-normalized data were projected to the reference to assign prediction scores and cell identities through anchor-based mapping. Predicted labels were reviewed and refined through expert inspection of canonical RNA and ADT marker expression, ensuring biologically coherent annotation across modalities. This approach identified 30 major immune populations encompassing lymphoid and myeloid lineages.

### Outlier Sample Removal

Following QC and initial uniform manifold approximation and projection (UMAP), a cluster of CD14+ monocytes was observed adjacent to the primary CD14+ monocyte cluster. This island consisted almost exclusively of three samples (two pediatric AD and one adult AD), suggesting a sample-specific artifact rather than a true biological subset. These outlier samples were excluded prior to Harmony^5^ integration, yielding a final dataset comprising 11 pediatric AD/HC, 13 adult AD/HC and 5 geriatric AD/HC participants.

### Data Integration

To correct for batch effects and align cells across sequencing runs, data integration was performed using Harmony implemented in Seurat v6. Principal component analysis (PCA) was conducted on SCTransform-normalized RNA expression, and the first 30 principal components were supplied to Harmony (RunHarmony, reduction = “pca”, dims = 1:30) with the *Batch* metadata variable as the grouping factor. The resulting Harmony embeddings were used to construct a shared nearest-neighbor (SNN) graph (FindNeighbors, reduction = “harmony”, dims = 1:30), identify clusters (FindClusters, resolution = 0.5), and generate UMAP visualizations (RunUMAP, reduction = “harmony”, dims = 1:30).

### Cell Type Proportion Analysis

Cell type frequencies were calculated at the subject level as the proportion of each annotated cell type relative to total numbers of singlet cells. For comparisons between AD and healthy controls, the Wilcoxon rank-sum test was applied. For three-group comparisons among pediatric, adult, and geriatric AD cohorts, both Kruskal–Wallis and pairwise Wilcoxon tests were performed. The Kruskal–Wallis test provided a global assessment of distributional differences across age groups, whereas pairwise Wilcoxon tests identified specific contrasts. Nominal *p*-values were reported, and cell-type proportions were visualized as grouped bar plots displaying mean ± standard error of the mean (SEM).

### Differential Gene and Protein Expression

Differential expression analyses were conducted separately for RNA (genes) and ADT (proteins) within each annotated cell type to identify molecular features associated with: 1) AD (all ages combined) vs healthy control (all ages combined) and 2) Between different AD age groups. Three intra-AD contrasts were tested independently: pediatric AD vs (adult AD + geriatric AD), adult AD vs (pediatric AD + geriatric AD), and geriatric AD vs (pediatric AD + adult AD). For RNA, differential expression was assessed using the Seurat FindMarkers function on the SCTransform-normalized assay with the Wilcoxon rank-sum test and significance thresholds of |log₂FC| ≥ 0.5 and false discovery rate (FDR) < 0.05. For proteins, differential expression used the CLR-normalized ADT assay with thresholds of log₂FC ≥ 0 and FDR < 0.05. No additional normalization was applied within subsets beyond global preprocessing. To visualize representative molecular programs across age groups, sentinel genes and proteins were selected from cell types with the highest numbers of significant DEGs and DEPs. For the most variable cell types, the top three features ranked by absolute log₂ fold-change were compiled. To ensure the resulting matrices captured biologically consistent and interpretable variation, features were retained only if they contained ≥ 2 non-missing (non-NA) log₂ fold-change values across (cell type × comparison) combinations and exhibited non-zero variance across comparisons.

### Machine Learning Model Development

Subject-level feature matrices were generated from the mean normalized expression of significant, differentially expressed genes and proteins. For RNA-based models, genes meeting significance thresholds (|log₂FC| ≥ 0.50 and FDR < 0.05) were selected from the intra-AD comparisons, and their SCTransform-normalized z-scores were averaged per subject within each cell type. For protein-based models, significant proteins (FDR < 0.05) were selected from ADT differential analyses, and CLR-normalized z-scores were averaged per subject within each cell type. These per-subject means were concatenated across all cell types to produce DEG- and DEP-based subject matrices. Subjects were stratified by age group and divided into 60 % training and 40 % validation partitions. To prevent data leakage, all preprocessing—including median imputation and scaling—was performed exclusively on the training set, and the resulting parameters were applied to the held-out validation data.

Feature discovery was performed using a random-forest one-vs-rest importance framework, in which MeanDecreaseAccuracy scores were computed across 5×3 repeated cross-validation folds within the training partition. Normalized importance scores were aggregated across folds, and the top 20 features were frozen for model evaluation. Model development employed nested internal cross-validation for hyperparameter tuning within the training set, followed by evaluation on the single 40% validation split. Eight supervised algorithms were implemented using the *caret* framework: random forest (rf), linear support vector machine (svmLinear), elastic net (glmnet), k-nearest neighbors (knn), naïve Bayes, linear discriminant analysis (lda), shallow neural networks (nnet), and averaged neural networks (avNNet). Hyperparameters were tuned to optimize predictive performance while constraining model complexity. For random forest, parameters included *mtry*, *ntree = 800*, and *nodesize = 3*; for linear SVM, the cost parameter *C* ranged from 0.01 to 10; for elastic net, *α* (0–0.5) and *λ* (1e⁻⁴–1) were varied; for k-NN, *k* values ranged from 5 to 11; for naïve Bayes, *laplace* (0–1) and *usekernel* (TRUE/FALSE) were tested; and for neural networks, hidden-layer size (1–2) and weight decay (0.01–0.1) were tuned with bagging disabled.

To address class imbalance and reduce variance, inverse-frequency class and case weights were applied according to model compatibility (e.g., *classwt* for random forest, *class.weights* for SVM, *weights* for neural networks). Overfitting was mitigated through model regularization, constrained tuning ranges, repeated cross-validation within the training partition, and the freezing of feature sets prior to validation. Model performance was evaluated using macro-averaged F1 as the primary metric, with area under the receiver operating characteristic curve (AUC) (one-vs-rest), overall accuracy, and Cohen’s κ as secondary measures. Random seeds (*set.seed = 742*) were fixed to ensure reproducibility. All analyses were conducted in R (v4.x) using *caret* and related packages.

## Results

### Clinical Characteristics of Subjects

After removing the sample outliers, our study cohort included a total of 58 participants spanning pediatric (0-17 years), adult (18-59 years), and geriatric (60+ years) age groups, each comprising both HC and individuals with AD (Supplementary Table 1, Additional File 1). The mean ages were 12.93 ± 4.84 (HC) and 12.36 ± 2.91 (AD) years for the pediatric group, 36.23 ± 13.23 and 36.00 ± 13.28 years for adults, and 70.60 ± 5.90 and 71.60 ± 5.94 years for the geriatric participants, respectively. Sex distribution was balanced across groups (overall 27 females and 31 males). The cohort was racially and ethnically diverse, with representation from White, Asian, Black/African American, Hispanic/Latino, and Native Hawaiian/Pacific Islander participants.

### Cell Types Enriched and Depleted in AD and Among Pediatric, Adult, and Geriatric AD

We characterized the immune cell composition in peripheral blood mononuclear cells (PBMCs) from 29 subjects with atopic dermatitis (AD) (11 pediatric, 13 adult, and 5 geriatric) and an equal number of matched healthy controls. The CITE-seq profiling captured high-quality singlets 282,197 total cells (median ≈ 4,445) with paired quantification of 140 surface proteins. The integrated dataset spanned 30 distinct immune subsets, encompassing major lymphoid and myeloid immune cell populations. These clusters were consistently represented across all six subgroups of pediatric, adult, and geriatric AD and HC, indicating robust integration and preservation of canonical PBMC identities (Figures 1A–B).

**Figure 1.**
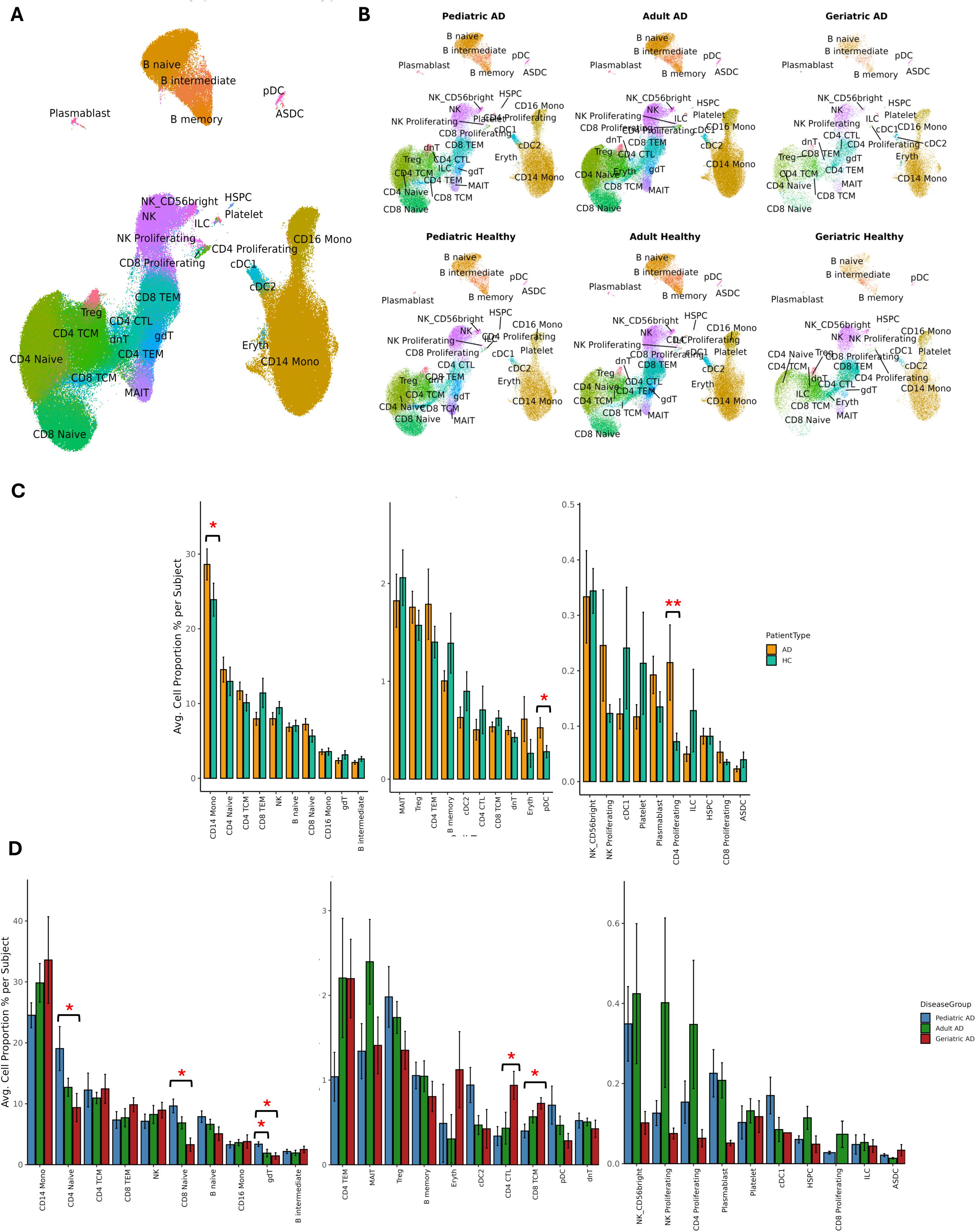
Immune landscape and cell-type composition in AD vs. HC and across AD age groups. (A) UMAP of all PBMCs showing 30 phenotypic subsets annotated by lineage and subset identity. (B) UMAPs split by patient status and age group (pediatric, adult, geriatric; AD and HC), demonstrating consistent representation of all major immune compartments across six subgroups. (C) Bar plots of mean cell-type proportions (AD vs HC) showing significant enrichment of CD14+ monocytes, pDCs, and CD4+ proliferating T cells (Wilcoxon, unadjusted p < 0.05). (D) Bar plots comparing AD subgroups only, highlighting significant age-linked differences in γδ T, CD4 naïve, and CD8 naïve populations (Kruskal–Wallis and pairwise Wilcoxon, p < 0.05).

When cell compositions were compared between AD and HC samples, CD14+ monocytes (p = 0.03) and plasmacytoid dendritic cells (pDC; p = 0.03) were significantly increased in AD, alongside an expansion of CD4+ proliferating T cells (p = 0.01) (Figure 1C). To investigate age effects within AD, we next restricted the analysis to AD patients and compared pediatric, adult, and geriatric subgroups (Figure 1D). Kruskal–Wallis’s statistics identified significant differences in γδ T cells (p = 0.01), CD8+ naïve T cells (p = 0.02), and CD4+ naïve T cells (p = 0.03). Pairwise analyses showed that pediatric AD exhibited higher cell-proportions of γδ T cells relative to both adult and geriatric groups (p = 0.01 for each comparison) and increased CD4+ and CD8+ naïve T cells compared to geriatric AD (p = 0.03 and 0.01, respectively). Conversely, CD4+ cytotoxic T cells (CD4+ CTL; p = 0.02) and CD8+ central memory T cells (CD8+ TCM; p = 0.04) were relatively elevated in geriatric AD. Together, these results reveal a developmental trajectory characterized by effector cell enrichment in AD with aging. These compositional trends prompted downstream transcriptomic and proteomic analyses to delineate the molecular programs underlying these age-linked immune shifts.

### Differentially Expressed Genes and Proteins Reveal Shared and Age-Specific Programs Across AD Age Groups

We next compared pediatric, adult, and geriatric AD by identifying differentially expressed genes (DEGs) and proteins (DEPs) across immune cell types. Individual subsets showed 220–467 DEGs (median = 310) and 3–18 DEPs (median = 6) distinguishing age groups, with the greatest transcriptomic changes in CD14+ monocytes, followed by CD4+ naïve, CD8+ naïve, B naïve, CD4+ TCM, and NK cells (Figure 2A). Surface protein differences were most pronounced in NK, CD4+ naïve, CD8+ naïve, B naïve, CD14+ monocyte, and mucosal-associated invariant T (MAIT) subsets (Figure 2B), reflecting partially overlapping but distinct age-associated patterns. To highlight representative signatures, the top three DEGs or DEPs per cell type and age-group contrast (AD vs. other AD groups) were selected, filtering out features with limited detection (fewer than two cell type–comparison combinations) or lacking variability in log₂ fold-change to ensure robust clustering across prioritized lineages (Figure 3A–3B).

**Figure 2.**
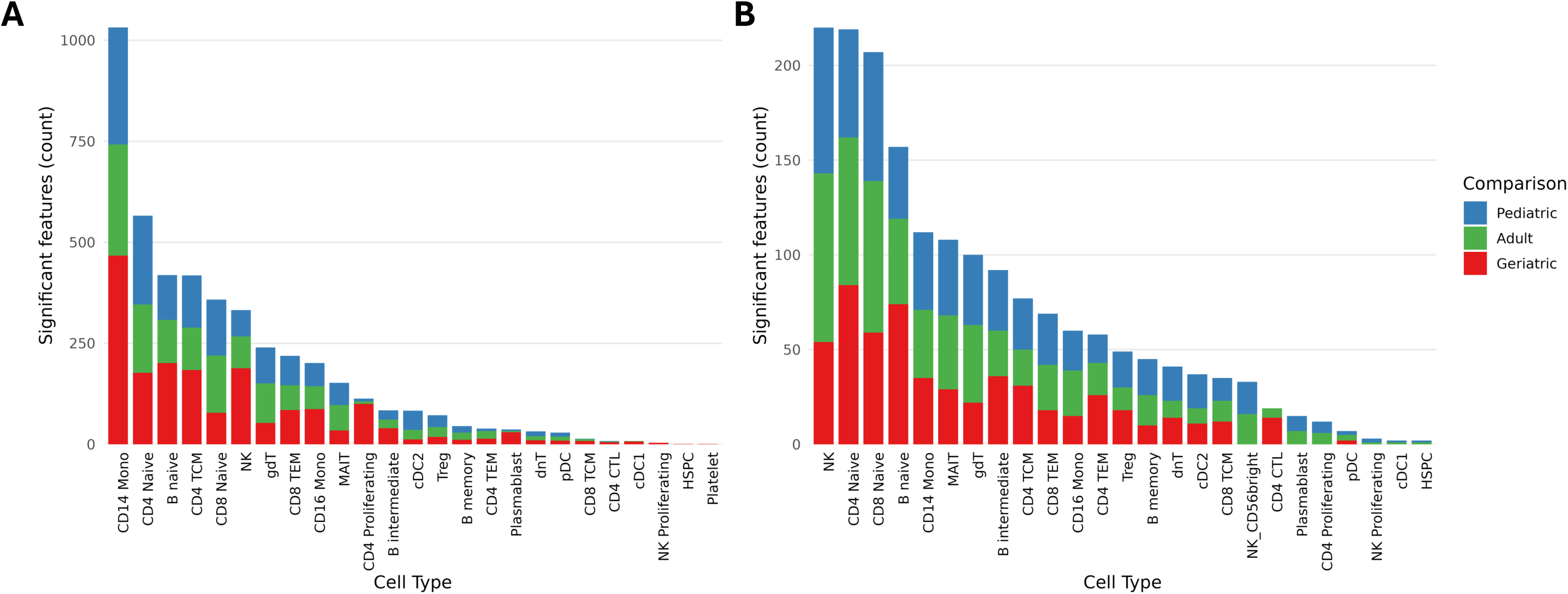
DEGs and DEPs by cell type and AD subgroup comparison. (A) Stacked bar plot of DEG counts per cell type × comparison (Pediatric AD vs Other AD, PvO, Adult AD vs Other AD, AvO, Geriatric AD vs Other AD, GvO). (B) Stacked bar plot of DEP counts per cell type × comparison.

**Figure 3.**
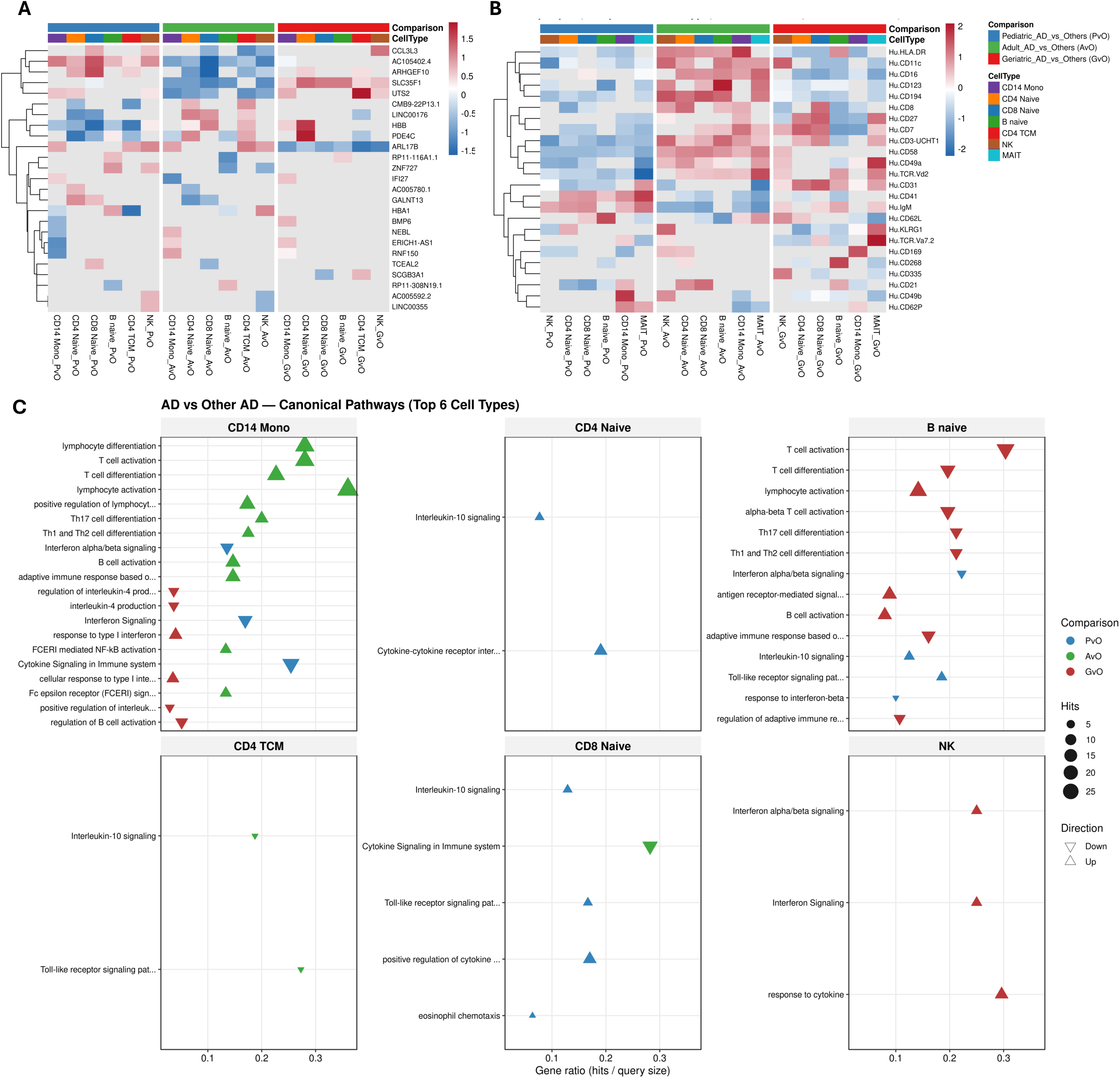
Transcriptomic and proteomic signatures reveal shared and age-specific programs across AD groups. (A) Heatmap of the top three “sentinel” differentially expressed genes (DEGs) per cell type and comparison—pediatric AD vs others (PvO), adult AD vs others (AvO), and geriatric AD vs others (GvO)—illustrating distinct transcriptional modules across CD14+ monocytes, CD4+ naïve, CD8+ naïve, B naïve, CD4+ TCM, and NK cells. (B) Heatmap of the top three “sentinel” differentially expressed proteins (DEPs) per cell type and comparison showing complementary remodeling of activation, adhesion, and immunoregulatory pathways across the same six lineages. (C) Pathway enrichment analysis of the full DEG set, highlighting canonical immune programs enriched by age group and cell type, including interferon/cytokine signaling, Th1/Th17 activation, chemotaxis, and cyclic adenosine monophosphate (cAMP) regulation. Collectively, these analyses reveal transcriptional and proteomic re-wiring across the AD lifespan—from interferon and cytokine activation in pediatric AD, to inflammatory and stress-responsive remodeling in adults, to metabolic and adhesion-driven adaptation in geriatric AD.

At the transcriptomic level, pediatric AD showed increased expression of transcription factor and signaling genes (e.g., *IFI27, ZNF727, ARHGEF10*) and trafficking or structural regulator *GALNT13*. The long noncoding RNA (lncRNA) *AC105402.4* was uniquely upregulated, suggesting age-specific epigenetic control. In adult AD, these same genes were downregulated, while differentiation and metabolic stress-related transcripts (e.g., *ERICH1-AS1, LINC00176, NEBL*) were elevated. Geriatric AD exhibited elevated signaling and metabolic regulators (e.g., *IFI27, BMP6, PDE4C, SLC35F1, HBB*) and persistent *ERICH1-AS1* expression, with unique downregulation of *ARL17B*.

Pathway analysis of the full DEG set (Figure 3C) showed enrichment of IL-10 and cytokine–cytokine receptor signaling in pediatric AD naïve compartments (B Naïve, CD4+ Naïve T, and CD8+ Naïve T). In terms of innate activation, toll-like receptor (TLR) activity was reduced specifically in CD14+ monocytes, but was upregulated in pediatric B Naïve and CD8+ Naïve T cells. In contrast, adult AD was enriched for monocyte-driven Th1/Th17/NF-κB activation and B-cell stimulation. Geriatric AD retained type-1 interferon activity and IL-10 signaling in monocytes and B cells but displayed reduced IL-4 production and T-cell signaling, while NK cells demonstrated elevated interferon and cytokine responses, reflecting a shift toward innate immune activation with aging.

Surface protein profiling revealed complementary remodeling across programs involving T-cell activation, trafficking, B-cell signaling, and innate regulation (Figure 3B). Pediatric AD showed upregulation of B-cell and immunoglobulin (B/Ig)-related proteins (e.g., IgM, CD41, CD62L) and downregulation of trafficking or naïve-lineage receptors (e.g., CD49a, CD8), suggesting early B-cell activity and a predominance of maturing lymphocytes. Adult AD exhibited stronger T-cell activation and differentiation, marked by increased activation, trafficking, and checkpoint proteins (HLA-DR, CD49a, CD21, KLRG1, CD16), consistent with enhanced effector and regulatory responses. Geriatric AD displayed upregulation of co-stimulatory and adhesion molecules (CD27, CD31, CD49a), together with myeloid and NK-associated markers (CD169, CD268, CD335) and unconventional T-cell receptors (TCR Vδ2, Vα7.2), reflecting adaptive and innate immune adjustments with aging.

### Machine Learning Classifiers Distinguish Pediatric, Adult, and Geriatric AD

We next evaluated the diagnostic potential of gene- and protein-level features to distinguish pediatric, adult, and geriatric AD. Using AD-only input matrices derived from significant DEGs and DEPs, we trained multi-class machine learning models to test whether compact molecular signatures could classify participants by age group. Model development used nested internal cross-validation for feature selection and hyperparameter tuning within the training set, followed by a single 40% held-out validation set for performance estimation. Due to the moderate sample size, particularly in the geriatric cohort, metrics reflect validation performance rather than full cross-subject cross-validation. Regularization and inverse-frequency class weighting were applied across all algorithms to control model complexity and mitigate class imbalance.

Across eight machine learning algorithms, DEG-based classifiers consistently outperformed DEP-based models. The avNNet and SVM Linear models yielded the highest validation performance, with avNNet reaching F1 = 0.70, AUC = 0.79, and accuracy = 0.64, while SVM Linear achieved AUC = 0.70 (Figure 4A). In contrast, DEP-based models demonstrated moderate discrimination with Naive Bayes performing best (macro F1 ≈ 0.63, AUC ≈ 0.71, accuracy ≈ 0.64) (Figure 4B). The geriatric AUC = 1.0 (Figure 5B) observed in the DEG validation set likely reflects data sparsity—only two geriatric samples in validation and three in training—rather than true perfect classification.

**Figure 4.**
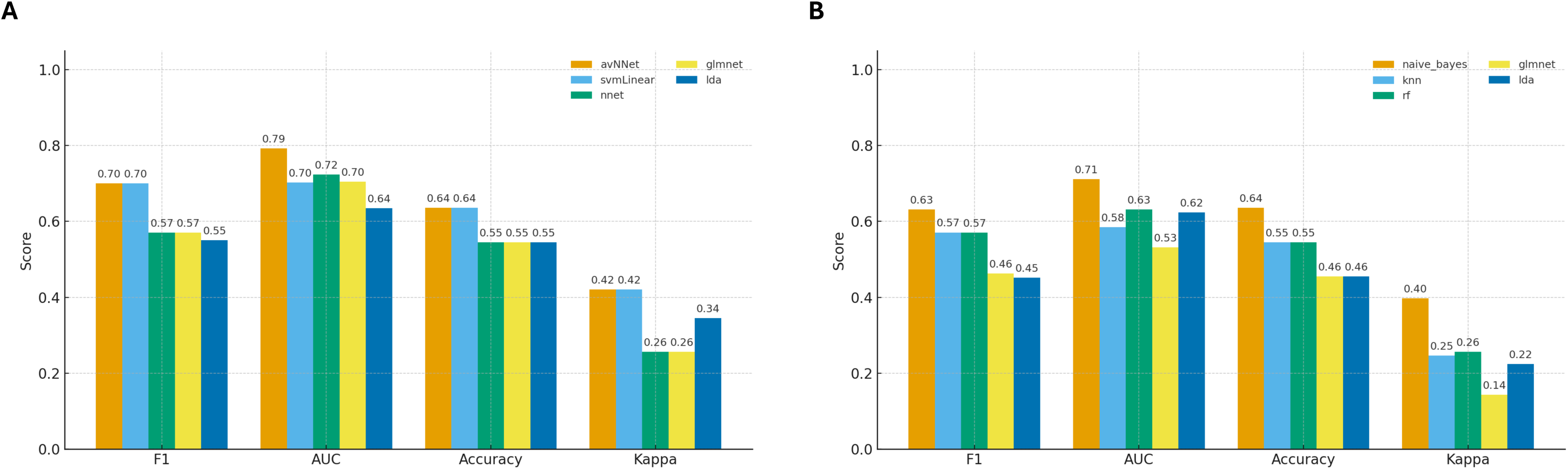
Performance of the top 5 machine learning classifiers. (A) Classifiers trained on DEG-based feature sets. (B) Classifiers trained on DEP-based feature sets.

**Figure 5.**
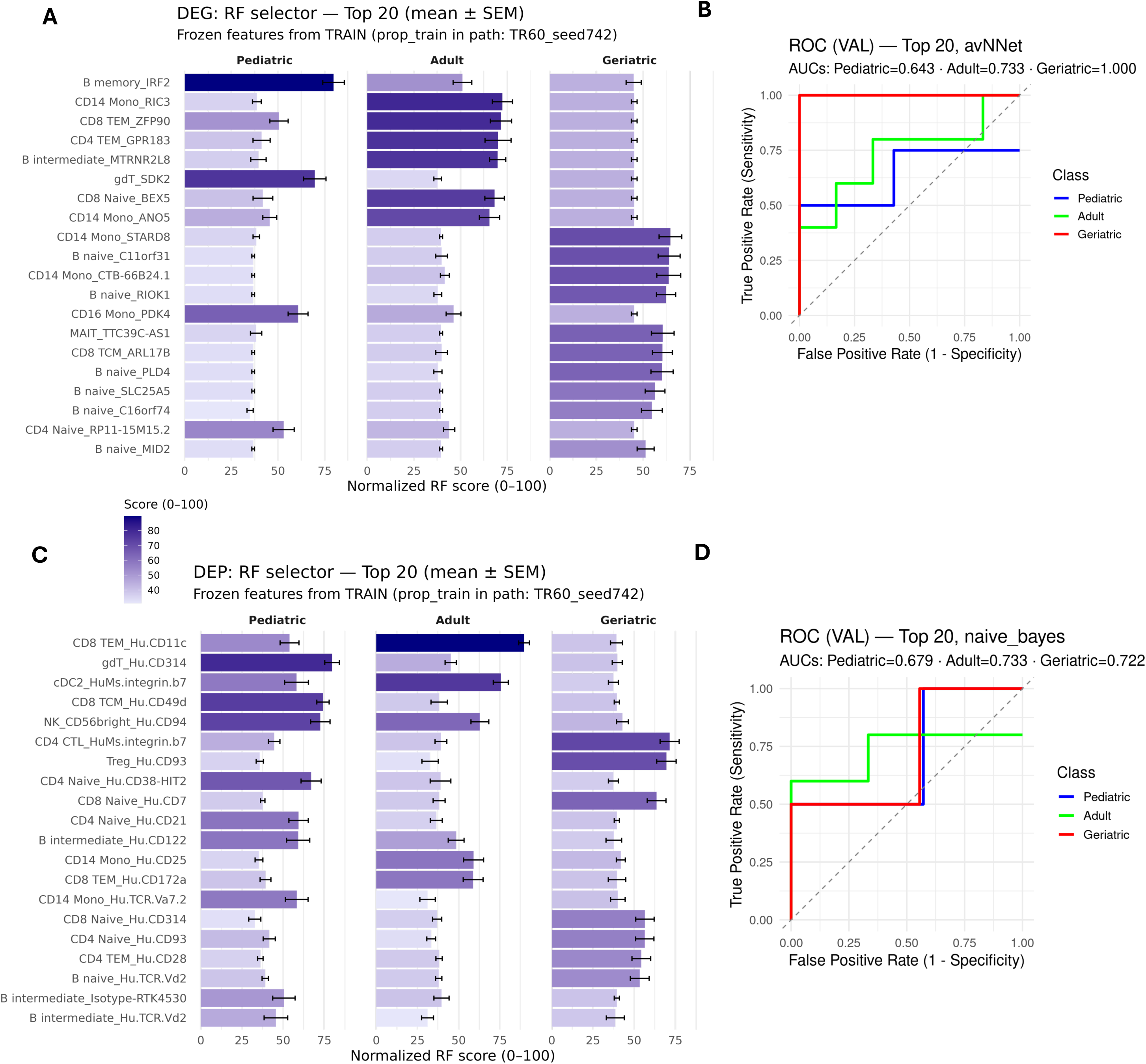
Feature importance across gene and protein modalities and validation model performance. (A) Top 20, stable gene features ranked by Random Forest importance. (B) Receiver operating characteristic (ROC) curves of the best-performing gene-based model (avNNet). (C) Top 20, stable protein features ranked by Random Forest importance. (D) ROC curves of the best-performing protein-based model (naïve Bayes).

The top 20 stable DEGs (Figure 5A) identified through Random Forest feature selection encompassed transcriptional and signaling regulators (*IRF2, ZFP90, GPR183, RIC3*), structural and trafficking mediators (*SDK2, STARD8, ANO5, MID2*), and epigenetic or lncRNA-associated genes (*CTB-66B24.1, RP11-15M15.2, ARL17B, TTC39C-AS1*). Additional features included protein synthesis and ubiquitin-related genes (*RIOK1, SLC25A5*) and metabolic regulators (*MTRNR2L8, PDK4*) (Figure 4B). In parallel, the top 20 DEPs identified through Random Forest feature selection (Figure 5C) encompassed T-cell activation and differentiation markers (CD28, CD25, CD38-HIT2), trafficking and homing receptors (CD49d, integrin β7, CD172a, CD11c), NK-related proteins (CD94, CD314), and B-cell or immunoglobulin-associated molecules (CD21, CD122). Additional γδ/MAIT T-cell receptors (TCR Vδ2, TCR Vα7.2) and regulatory proteins (CD93) reflected the integrated remodeling of adaptive and innate immune programs across age groups.

Collectively, these results demonstrate that compact, biologically interpretable feature sets derived from differential expression are sufficient to separate AD age groups with moderate-to-high accuracy, supporting their potential use as stable diagnostic signatures for downstream biological interpretation.

### Biological Interpretation of Stable Gene and Protein Features Across AD Age Groups

We next examined the biological relevance of these stable DEGs and DEPs. These features collectively captured transcriptional, structural, and metabolic programs that differentiated pediatric, adult, and geriatric AD while maintaining cell-type diversity across B cells, monocytes, and T-cell lineages (Figure 6A–B). The Z-scored expression of the stable top 20 gene and protein features is detailed in heatmaps in Additional File 3, illustrating the consistent, age-dependent transcriptional and proteomic remodeling that we subsequently analyzed by functional group (Supplementary Tables 3-4, Additional Files 4-5).

**Figure 6.**
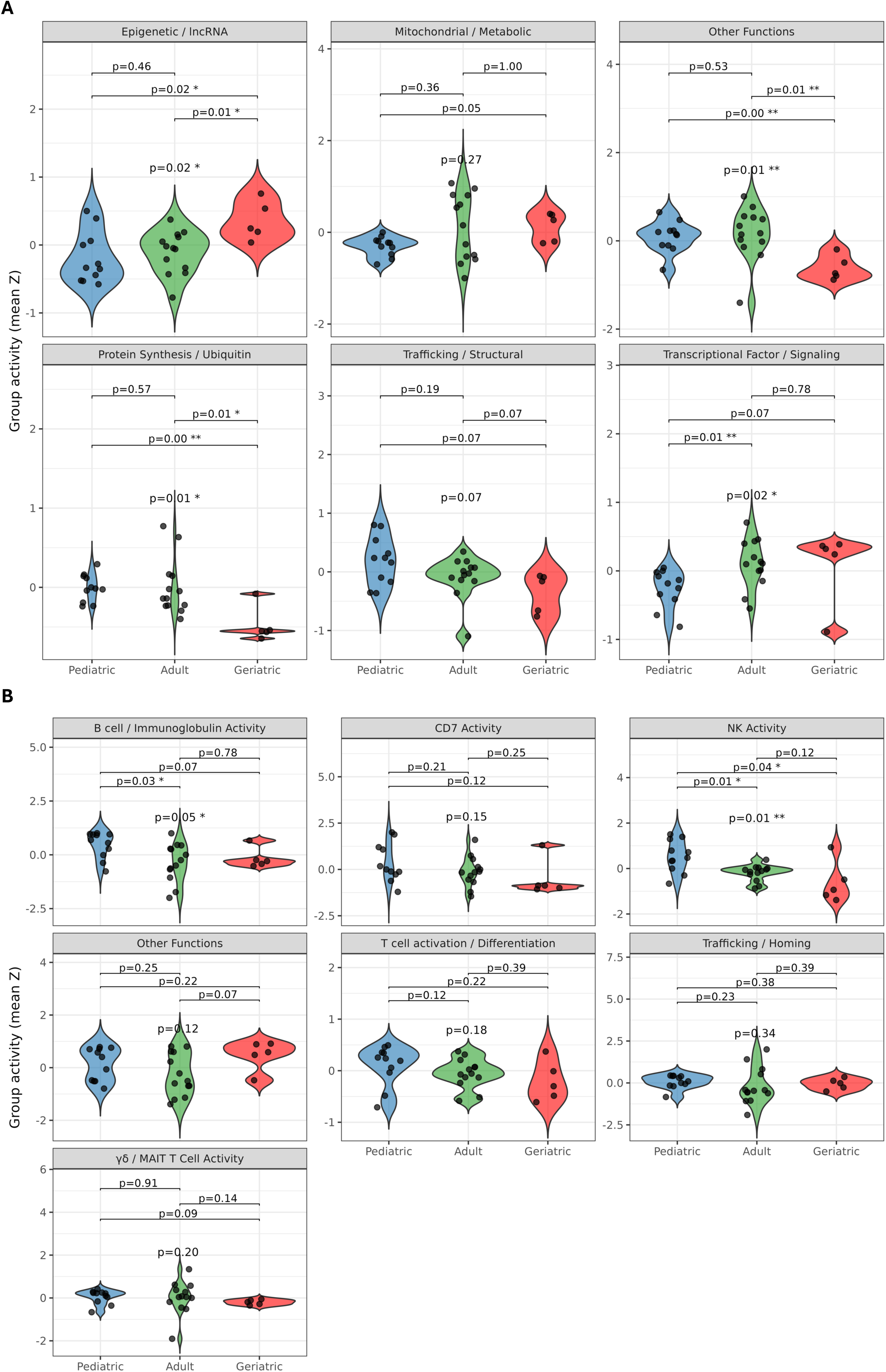
Functional group activity across stable genes and proteins. (A) Violin plots showing mean Z-scored expression of the top 20 stable genes grouped by functional activity. (B) Violin plots showing mean Z-scored expression of the top 20, stable proteins grouped by functional activity.

Significant age-associated variation was observed in transcriptional signaling, protein synthesis/ubiquitin, and epigenetic/lncRNA regulation (p = 0.02, 0.01, and 0.02, respectively; Figure 6A). While pediatric AD exhibited modest, non-significant elevations in structural and trafficking genes (e.g., *SDK2, STARD8*), these trends suggest early cytoskeletal and translational readiness rather than full immune activation. In contrast, adult AD demonstrated pronounced upregulation of transcriptional regulators (*IRF2, ZFP90*) and metabolic mediators (*PDK4, RIC3*) compared to pediatric AD (p = 0.01), indicating peak transcriptional and metabolic remodeling characteristic of chronic inflammation. Geriatric AD, by comparison, displayed significantly reduced expression of protein synthesis/ubiquitin (*RIOK1, SLC25A5*; p = 0.01 compared to pediatric AD and p = 0.01 compared to adult AD) and lncRNA-associated genes (*RP11-15M15.2*; p = 0.02 compared to pediatric AD and p = 0.01 compared to adult AD), indicating diminished translational activity and regulatory control consistent with a compensatory shift from adaptive to innate immunity during inflammaging. Collectively, these findings reveal a trajectory in which adult AD represents the inflammatory apex, flanked by pediatric AD’s early structural activation and geriatric AD’s adaptive-to-innate transition.

At the protein level, significant age-associated differences were observed in NK Activity (p = 0.01) and B cell/Immunoglobulin activity (p = 0.05) (Figure 6B). Both programs showed the highest expression in pediatric AD, with lower levels in adult (p = 0.01, 0.03, respectively) and geriatric (p = 0.04) groups, indicating stronger early NK- and B-cell engagement that diminishes with age. Key proteins driving these differences included NK-related CD94 and CD314 and B/Ig-related CD21 and CD122. In contrast, T-cell activation and differentiation (CD28, CD25, CD38-HIT2), trafficking and homing (CD49d, integrin β7, CD172a, CD11c), and γδ/MAIT T-cell receptors (TCR Vδ2, TCR Vα7.2) remained relatively stable across age groups, reflecting preserved effector and adhesion programs despite chronic inflammation. Together, these findings highlight that pediatric AD is characterized by amplified NK and B-cell activation, whereas adult and geriatric AD display attenuated activity within these arms.

## Discussion

Our study sheds light on immune composition differences among pediatric, adult, and geriatric patients with AD across multiple levels of resolution from PBMC. At the cell composition level, AD patients showed higher proportions of CD14+ monocytes and pDC cells compared with healthy controls, consistent with earlier reports of altered dendritic-cell balance in AD ^6,7^. Within the AD cohort, pediatric patients had more γδ T cells and naïve CD4+ and CD8+ T cells than adults or older patients, suggesting a less mature immune profile with greater naïve T-cell proportions early in life^8^. In contrast, geriatric AD was marked by an increase in CD4+ CTL and CD8+ TCM, reflecting a shift toward more mature and effector cell types. Together, these changes suggest that AD follows the general pattern of immune aging, characterized by reduced naïve and increased effector and memory populations ^9^, but exacerbates it ^10^. Our subsequent analyses provide a potential mechanism for this exacerbation, defining geriatric AD as a ‘hybrid inflammaging’ state. This state is measurably distinct from pediatric and adult AD groups and is characterized by a unique shift toward enhanced innate signaling, compensatory regulatory circuits, and specific myeloid protein expression.

Within the most variable immune cell types (CD14+ monocytes, CD4+ naïve, CD8+ naïve, B naïve, CD4+ CTM, and NK cells), we observed progressive age-linked remodeling of immune programs in AD (Figure 1B). In pediatric AD, upregulation of transcriptional and signaling regulators (*IFI27, ZNF727, ARHGEF10*) and the structural gene (*GALNT13*), together with the lncRNA *AC105402.4*, indicates coordinated activation of regulatory and structural pathways that overlap with developmental and differentiation programs. This pattern, accompanied by enriched IL-10 and cytokine–cytokine receptor signaling, suggests that early immune programs remain active, potentially reflecting sustained tissue–immune remodeling under chronic inflammatory stress. Because IL-10, an anti-inflammatory cytokine that dampens the immune system’s overreaction to allergens, was enriched in pediatric AD, this enrichment may explain the absence of the canonical Th2 skewing typically reported in this age group ^11,12^. Th2-driven inflammation, which characterizes allergen-associated immune activation in AD, was instead more pronounced in adult AD, particularly within CD14+ monocytes. The elevated IL-10 activity observed in pediatric naïve T- and B-cell compartments may have restrained Th2 polarization, consistent with IL-10’s established role in allergen tolerance and immune regulation^13^. Persistent IL-10 signaling under chronic inflammatory stress may therefore prolong developmental tolerance programs, maintaining regulatory circuits beyond their typical maturation window. The concurrent reduction of toll-like receptor signaling in CD14+ monocytes but increased responsiveness in naïve B and T cells supports selective restraint of innate pathways alongside heightened adaptive priming, consistent with early-life immune plasticity ^14^. In adult AD, the downregulation of early transcriptional regulators (*IFI27, ZNF727, ARHGEF10*) and upregulation of mitochondrial, metabolic, and stress-response genes (*PDK4, ERICH1-AS1, LINC00176, NEBL*) mark a shift from developmental plasticity toward inflammation-driven mitochondrial and metabolic reprogramming^15^. Pathway enrichment for Th2, Th1, Th17, and NF-κB signaling, particularly within CD14+ monocytes, indicates broad effector remodeling consistent with reports that adult AD exhibits stronger Th1/Th17 activity than pediatric AD^12^. The concurrent decrease of IL-10 and TLR signaling in CD4+ central-memory T cells reflects diminishment of the tolerance network and the emergence of an effector-dominant inflammatory state. In geriatric AD, persistent interferon and IL-10 signaling accompanied higher expression of metabolic and mitochondrial regulators (*BMP6, PDE4C, SLC35F1, HBB*) and retention of *ERICH1-AS1* expression, suggesting adaptive, energy-conserving responses typical of immune aging^9,15^. Reduced IL-4 production and T-cell signaling, together with enhanced NK-cell interferon and cytokine activity, further highlight a shift toward innate immune compensation. The presence of adhesion and co-stimulatory genes (*BMP6, SLC35F1*) and increased myeloid-associated proteins (CD169, CD268, CD335) reinforces a hybrid “inflammaging” state—defined by chronic low-grade inflammation coupled with compensatory regulatory signaling within the aging immune network ^7^.

Surface-protein analyses supported these transcriptomic trends. In pediatric AD, upregulation of B-cell and immunoglobulin-related proteins (IgM, CD41, CD62L) and downregulation of trafficking and γδ T-cell receptor markers (CD49a, TCR Vδ2) indicate early B-cell activity and a predominance of maturing lymphocytes. While enrichment of naïve and transitional B-cell subsets has been reported in adult AD blood^12^, our findings suggest that humoral activation may arise earlier in disease development, already detectable in pediatric AD. In adult AD, elevated expression of activation, trafficking, and checkpoint proteins marks the peak of chronic immune activation. Increased antigen presentation (HLA-DR), T-cell trafficking (CD49a), B-cell activation (CD21), and cytotoxic and effector activity (KLRG1, CD16) reflect heightened engagement of both adaptive and innate pathways characteristic of sustained inflammatory remodeling ^8,11,7^. This profile aligns with our transcriptomic findings of enhanced Th1/Th17 and NF-kB signaling, reinforcing that adult AD represents a state of maximal immune activation and metabolic stress. In geriatric AD, upregulation of co-stimulatory and adhesion molecules (CD27, CD31), myeloid and NK-associated proteins (CD169, CD268, CD335), and unconventional T-cell receptors (TCR Vδ2, Vα7.2) reflects a shift toward enhanced innate signaling and reduced adaptive dominance^7^. Notably, CD27 expression was increased in both adult and geriatric AD, consistent with previous reports in adult AD cohorts^11^. In adults, sustained CD27 expression likely reflects ongoing adaptive activation through NF-kB-mediated survival and memory signaling^16^, whereas in geriatrics, elevated CD27 may represent compensatory co-stimulatory signaling amid declining adaptive function^9^. This interpretation aligns with reports of reduced NK cytotoxicity, heightened monocyte activation, and expansion of adhesion and co-stimulatory networks with aging AD^7^.

Machine learning–based feature selection and average expression analyses revealed compact, biologically coherent gene and protein signatures that distinguish pediatric, adult, and geriatric AD with moderate-to-high accuracy. Transcript-level classifiers (avNNet, SVM Linear) achieved superior performance (F1 ≈ 0.70) compared to protein-based classifiers (F1 ≈ 0.63), reflecting greater transcriptomic stability across individuals. In pediatric AD, key ML features such as CD21, CD94, and CD122 reaffirmed the early B-cell and NK-cell activation observed in the differential protein analysis, supporting a state of heightened humoral and innate readiness rather than full inflammatory activation. In adult AD, the strong weighting of RIC3, ZFP90, GPR183, MTRNR2L8, BEX5, and ANO5 confirmed peak transcriptional and metabolic remodeling, consistent with the DEG findings, which capture the convergence of adaptive and metabolic stress responses typical of chronic inflammation. In geriatric AD, the prominence of co-stimulatory, adhesion, and innate activation markers mirrored the DEP findings, reflecting a shift toward innate signaling and compensatory immune activation characteristic of inflammaging. When aligned with the transcriptomic and surface-protein analyses, these ML-derived features delineate a coherent age trajectory: pediatric AD marked by early innate and humoral priming, adult AD by maximal transcriptional and metabolic activation, and geriatric AD by adaptive contraction and innate compensation. Together, these patterns demonstrate that compact, cell-informed molecular signatures can capture the immunologic trends from developmental immune readiness to chronic inflammatory remodeling and age-related decline in AD.

There were several limitations to this study. Sample collection was cross-sectional and did not track patients longitudinally, preventing direct assessment of temporal immune shifts. However, we minimized confounding by excluding subjects receiving systemic immunomodulators at the time of sampling. The moderate overall sample size, particularly within the geriatric group, limited model generalizability and constrained feature selection in machine-learning analyses. Additionally, our data were restricted to circulating immune cells, which may not fully capture skin-resident or tissue-specific activity. Nonetheless, the concordant transcriptomic and proteomic trends across immune lineages, supported by internal cross-validation, strengthen the robustness of these findings. Future studies integrating matched skin–blood single-cell data, longitudinal sampling, and perturbation experiments will be essential to validate these pathways and evaluate therapeutic relevance.

## Conclusion

Through single-cell multi-omics profiling, we identified distinct immunologic differences across the atopic dermatitis lifespan. These age-linked immune programs yielded stable molecular signatures capable of distinguishing AD subgroups, providing a foundation for age-specific diagnostics, prognostic refinement, and tailored therapeutic strategies for atopic dermatitis across the human lifespan.

## Abbreviations

- AD: Atopic dermatitis
- HC: Healthy control
- PBMCs: Peripheral blood mononuclear cells
- CITE-seq: Cellular Indexing of Transcriptomes and Epitopes by sequencing
- RNA: Ribonucleic acid
- ADT: Antibody-derived tag
- vIGA-AD: Validated Investigator Global Assessment for Atopic Dermatitis
- IRB: Institutional Review Board
- ACD: Acid citrate dextrose
- FBS: Fetal bovine serum
- DMSO: Dimethyl sulfoxide
- SNP: Single nucleotide polymorphism
- VCF: Variant call format
- GRCh37/GRCh38: Genome Reference Consortium Human build 37/38
- QC: Quality control
- UMAP: Uniform Manifold Approximation and Projection
- PCA: Principal component analysis
- SNN: Shared nearest neighbor
- DEG(s): Differentially expressed gene(s)
- DEP(s): Differentially expressed protein(s)
- FDR: False discovery rate
- SEM: Standard error of the mean
- CTL: Cytotoxic T lymphocyte
- TCM: Central memory T cell
- NK: Natural killer
- MAIT: Mucosal-associated invariant T cell
- Th1/Th2/Th17/Th22: T helper 1/2/17/22
- JAK–STAT: Janus kinase–signal transducer and activator of transcription
- TLR: Toll-like receptor
- NF-κB: Nuclear factor kappa-light-chain-enhancer of activated B cells
- cAMP: Cyclic adenosine monophosphate
- AUC: Area under the receiver operating characteristic curve
- AUROC: Area under the receiver operating characteristic curve (multi-class context)
- ROC: Receiver operating characteristic
- lncRNA: Long noncoding RNA

## Declaration

### Ethics approval and consent to participate

All participants provided written informed consent under UCSF Institutional Review Board approval 10-02830 and in accordance with the Declaration of Helsinki rules.

### Consent for publication

Not applicable.

### Availability of data and materials

The data supporting this study’s conclusions are included within the article and its supplementary materials. Additional datasets are available from the corresponding author upon reasonable request.

### Competing interests

W.L. has received research grants from Amgen, Janssen, Leo, and Regeneron. T.B. has received research funding from Amgen, Castle, CorEvitas, Novartis, Pfizer, and Regeneron. She has served as an advisor for Abbvie, Apogee, Arcutis, Aslan, Boehringer-Ingelheim, Bristol Myers Squibb, Castle, Dermavant, Galderma, Incyte, Janssen, Leo, Lilly, Oruka, Pfizer, Novartis, Sanofi, Sun, Takeda, Taxa, and UCB. She is a speaker for Abbvie, Amgen, Arcutis, BMS, Dermavant, Galderma, Janssen, Lilly, Leo, Mindera, Ortho, Sanofi, and UCB. E.M. has received consulting fees from Novartis. The other authors report no conflicts of interest in this work.

### Funding

The work was funded by grant NEA21-ECRG158 from the National Eczema Association to W.L. W.L.is also supported by grants UC2AR081029, R01AR078688, and R21AR084041.

### Authors’ contributions

Conceptualization (W.L., G.C.B.), Methodology (W.L., G.C.B., S.K.), Software (G.C.B., S.K.),

Validation (G.C.B., S.K.), Formal analysis (G.C.B.), Investigation (G.C.B., J.J., A.I., M.B., I.N.,

E.M., I.N.), Resources (W.L., S.K., X.F.), Data Curation (S.K., G.C.B.), Writing – Original Draft

(G.C.B.), Writing – Review & Editing (G.C.B., W.L., S.K.), Visualization (G.C.B.), Project Administration (W.L., G.C.B., S.K.), Funding Acquisition (W.L.)

## Acknowledgement

Gian Carlo Baldonado would like to thank Wilson Liao for their administrative support, mentorship, and research guidance, as well as Sugandh Kumar for their mentorship and technical guidance. We would like to thank Bobby Shih and the fellows and medical students from the UCSF Psoriasis Center for their contributions.

## Authors’ information (optional)

Not applicable.

## Additional Files

### Additional File 1

- File format: .xls
- Title of data: Supplementary Table 1. Subject clinical and demographic characteristics.

### Additional File 2

- File format: .xls
- Title of data: Supplementary Table 2. 140-plex TotalSeq-B antibody panel.
- Description of data: List of 140 antibodies included in the TotalSeq-B panel used for cell staining.

### Additional File 3

- File format: .pdf
- Title of data: Supplementary Figure 1. Z-scored average expression of top stable gene and protein features.
- Description of data: Heatmaps showing Z-scored average expression for (A) the top 20, stable genes and (B) the top 20, stable proteins.

### Additional File 4

- File format: .xls
- Title of data: Supplementary Table 3. Transcriptomic functional groups.
- Description of data: Mapping of top stable genes to their functional groups used in Figure 6

### Additional File 5

- File format: .xls
- Title of data: Supplementary Table 3. Proteomic functional groups.
- Description of data: Mapping of top stable proteins to their functional groups used in Figure 6.

## References

1. Ständer S. Atopic Dermatitis. Ropper AH, ed. N Engl J Med. 2021;384(12):1136-1143. doi:10.1056/NEJMra2023911

2. Bieber T. Atopic dermatitis: an expanding therapeutic pipeline for a complex disease. Nat Rev Drug Discov. 2022;21(1):21-40. doi:10.1038/s41573-021-00266-6

3. Liu J, Kumar S, Hong J, et al. Combined Single Cell Transcriptome and Surface Epitope Profiling Identifies Potential Biomarkers of Psoriatic Arthritis and Facilitates Diagnosis via Machine Learning. Front Immunol. 2022;13:835760. doi:10.3389/fimmu.2022.835760

4. Kang HM, Subramaniam M, Targ S, et al. Multiplexed droplet single-cell RNA-sequencing using natural genetic variation. Nat Biotechnol. 2018;36(1):89-94. doi:10.1038/nbt.4042

5. Korsunsky I, Millard N, Fan J, et al. Fast, sensitive and accurate integration of single-cell data with Harmony. Nat Methods. 2019;16(12):1289-1296. doi:10.1038/s41592-019-0619-0

6. Dias de Oliveira NF, Santi CG, Maruta CW, Aoki V. Plasmacytoid dendritic cells in dermatology. An Bras Dermatol. 2021;96(1):76-81. doi:10.1016/j.abd.2020.08.006

7. Jin S, Lee K, Bang YJ, et al. Mapping the immune cell landscape of severe atopic dermatitis by single-cell RNA -seq. Allergy. 2024;79(6):1584-1597. doi:10.1111/all.16121

8. Esaki H, Czarnowicki T, Gonzalez J, et al. Accelerated T-cell activation and differentiation of polar subsets characterizes early atopic dermatitis development. J Allergy Clin Immunol. 2016;138(5):1473-1477.e5. doi:10.1016/j.jaci.2016.04.052

9. Fulop T, Larbi A, Dupuis G, et al. Immunosenescence and Inflamm-Aging As Two Sides of the Same Coin: Friends or Foes? Front Immunol. 2018;8:1960. doi:10.3389/fimmu.2017.01960

10. Chen PY, Shen M, Cai SQ, Tang ZW. Association Between Atopic Dermatitis and Aging: Clinical Observations and Underlying Mechanisms. J Inflamm Res. 2024;17:3433-3448. doi:10.2147/JIR.S467099

11. Czarnowicki T, Gonzalez J, Bonifacio KM, et al. Diverse activation and differentiation of multiple B-cell subsets in patients with atopic dermatitis but not in patients with psoriasis. J Allergy Clin Immunol. 2016;137(1):118-129.e5. doi:10.1016/j.jaci.2015.08.027

12. Jeong S, Shin S, Choi HS, et al. Pediatric Atopic Dermatitis Exhibits Distinctive Patterns in JAK/STAT Pathway Activation and CD6-ALCAM Signaling. Immune Netw. 2025;25(4):e26. doi:10.4110/in.2025.25.e26

13. Carlini V, Noonan DM, Abdalalem E, et al. The multifaceted nature of IL-10: regulation, role in immunological homeostasis and its relevance to cancer, COVID-19 and post-COVID conditions. Front Immunol. 2023;14:1161067. doi:10.3389/fimmu.2023.1161067

14. Pieren DKJ, Boer MC, De Wit J. The adaptive immune system in early life: The shift makes it count. Front Immunol. 2022;13:1031924. doi:10.3389/fimmu.2022.1031924

15. Chen Y, Ye Y, Krauß PL, et al. Age-related increase of mitochondrial content in human memory CD4+ T cells contributes to ROS-mediated increased expression of proinflammatory cytokines. Front Immunol. 2022;13:911050. doi:10.3389/fimmu.2022.911050

16. Song XF, Lin F, Chen ZG, Zhao GA, Sun SY, Pu J. The Relationship and Mechanism of CD27 with Coronary Atherosclerotic Heart Disease. Cardiovasc Drugs Ther. Published online August 8, 2025. doi:10.1007/s10557-025-07740-y

