## Supplementary figures and images for "Immunological Differences in Atopic Dermatitis Across Age Groups: Insights from Single-Cell Multi-Omics"

### Supplementary Figure 1

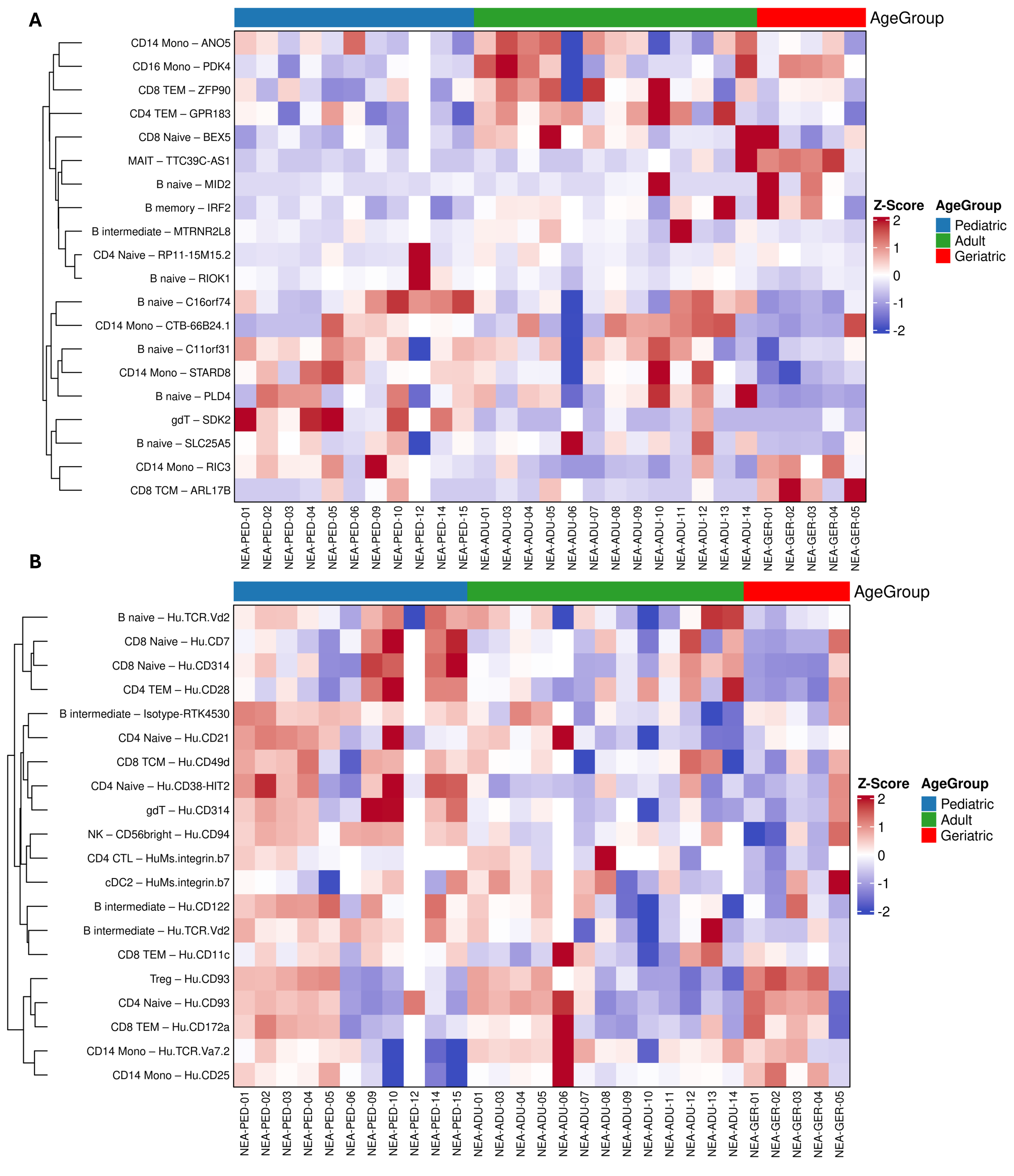
